# The Impact of Preprints on COVID-19 Research Dissemination: A Quantitative Analysis of Journal Publications

**DOI:** 10.1101/2024.05.28.596359

**Authors:** Hiroyuki Tsunoda, Yuan Sun, Masaki Nishizawa, Xiaomin Liu, Kou Amano, Rie Kominami

## Abstract

Preprints have played an unprecedented role in disseminating COVID-19-related science results to the public. The study aims to elucidate the role of preprints during the COVID-19 public health emergency (2020-2023) through a quantitative analysis of journal papers. Among the 247,854 COVID-19-related papers published in PubMed, 12,152 were initially released as preprints and were eventually published in 1,380 journals. This number is more than five times the 246 journals to which submissions can be made directly from bioRxiv through the B2J program. Journals with higher impact factors and Normalized Eigenfactor scores tend to publish a larger number of preprint-derived articles. The proportion of preprints among PubMed papers was 0.049, but this varies significantly by journal. In the top 30 journals, most exceed this proportion, indicating that these journals are preferred by authors for submitting their work. These findings highlight the growing acceptance and impact of preprints in the scientific community, particularly in high-impact journals.

## Introduction

Preprints are open and accessible scientific manuscripts or reports that are shared publicly through preprint archives before being submitted to journals. The value and importance of preprints have grown since their contribution during the public health emergency of the COVID-19 pandemic (Blatch-Jones, et al., 2023). Some researchers argue that because preprints are not peer-reviewed, the quality of the research cannot be guaranteed. Preprints were not very characteristic of citation practices in many disciplines (Tenopir, et al., 2016). However, as of October 2022, half of the preprints related to COVID-19 posted on medRxiv in 2020 were later published in peer-reviewed journals (Llor, Moragas, & Maier, 2022). Moreover, many of the peer-reviewed journals that published drafts derived from preprints were part of the Web of Science Core Collection by Clarivate Analytics and had high Journal Impact Factors (Tsunoda, et al., 2019). The quality of reporting in preprints in the life sciences is within a similar range as that of peer-reviewed papers, albeit slightly lower on average, supporting the idea that preprints should be considered valid scientific contributions (Carneiro, et al., 2020; Davidson, et al., 2024; Janda, et al., 2022; Sarkis-Onofre, Girotto, & Agostini, 2023). Research papers initially disseminated as preprints tend to undergo a shorter peer review process when submitted to journals, leading to faster publication (Fraser, Mayr, & Peters, 2022; Tsunoda, et al., 2020). Moreover, studies have shown that the peer review duration for papers shared as preprints is further reduced if authors opt to publish them as preprints before submitting them to journals (Tsunoda, et al., 2022). Preprints have proliferated since the COVID-19 pandemic, and the number of journal papers initially disseminated as preprints has also increased (Gianola, et al., 2020). Preprints indicate that papers related to COVID-19 are published to journals at a higher frequency than those unrelated to COVID-19 (Otridge et al., 2022; Tsunoda et al., 2023b) and undergo peer review more expeditiously (Fraser, et al., 2021; Tsunoda, et al., 2023a). Therefore, publishing research results in preprints can quickly communicate research results even to researchers who do not use preprints. Preprints have played an unprecedented role in disseminating COVID-19-related science results to the public (Nelson, et al., 2022; Wang, et al., 2021; Zeng, 2023). The study aims to elucidate preprints’ role during the COVID-19 public health emergency (2020-2023) through quantitative analysis of journal papers.

## Data and Methods

The materials analyzed in this study consist of papers, henceforth referred to as PubMed papers, which were published in journals between January 2019 and December 2023. These papers were registered in PubMed and assigned the COVID-19 MeSH (Medical Subject Headings) term. Data retrieval was conducted using the public API NCBI Entrez Programming Utilities between April and May 2024, resulting in the collection of 247,854 papers. Duplicate entries were removed using PubMed IDs as unique keys. The preprints were restricted to papers published on two platforms, bioRxiv and medRxiv. Papers initially disseminated as preprints were defined as those submitted to and subsequently published in a journal following their initial publication as preprints. Preprints published on bioRxiv or medRxiv during this period were downloaded using the bioRxiv API, and duplicates were removed using DOIs as keys, resulting in 126,164 preprints. After screening, we combined PubMed articles using the DOI as a key and found 12,152 preprint matches, removing 114,016 preprints.

The analysis involved combining the set of 247,854 PubMed articles, including the 12,152 preprint-derived articles, to determine the proportion of COVID-19-related preprints among the COVID-19-related PubMed papers in journals. The Journal Impact Factor (JIF) 2022 and Normalized Eigenfactor were obtained from the Journal Citation Reports (JCR) 2022 created by Clarivate. We combined the two sets of 1,380 journals and JCR2022 using ISSNs as the key, resulting in a total of 1,249 journals with assigned JIF and Normalized Eigenfactor scores. Spearman’s rank correlation coefficients were calculated to measure the relationship between the number of preprints and JIF, the number of preprints and Normalized Eigenfactor, and the proportion of preprints and JIF.

## Results

Among the 247,854 COVID-19-related papers published in PubMed, 12,152 were initially released as preprints and were eventually published in 1,380 journals. This number is more than five times the 246 journals to which submissions can be made directly from bioRxiv through the B2J (DIRECT TRANSFER FROM BIORXIV TO JOURNALS OR PEER REVIEW SERVICES) program. This discrepancy highlights the broad acceptance and integration of preprints across a wide range of journals. It suggests that preprints are becoming an essential part of the scientific communication process, reaching a diverse array of journals and audiences.

The analysis also revealed significant correlations between the number of preprints in journals and journal metrics. The Spearman’s rank correlation coefficients were 0.3762 between preprints and Journal Impact Factor (JIF), 0.4442 between preprints and Normalized Eigenfactor, and 0.0986 between the proportion of preprints and JIF, all indicating a p-value of less than 1%, demonstrating significance.

The proportion of preprints among PubMed papers was 0.049. However, this situation varies significantly by journal. In the top 30 journals, most exceed this proportion, indicating that these are preferred journals for authors to submit their work. Notably, these 30 journals accounted for approximately 50% of all preprint-derived articles published in the 1,380 PubMed journals. High JIF journals such as *Nature* (64.8), *Cell* (64.5), *Science* (56.9), *Eurosurveillance* (19.0), *Nature Communications* (16.6), and *JAMA Network Open* (13.8), as well as journals with high Normalized Eigenfactor scores such as *Nature Communications* (303.23), *Science* (237.43), *Scientific Reports* (227.88), *PLOS ONE* (153.58), and *Proceedings of the National Academy of Sciences of the United States of America* (146.31), were ranked among the top 30 in terms of the number of preprints.

**Table 1.** Top 30 journals frequently publishing COVID-19-related papers initially disseminated as preprints. [JIF: Journal Impact Factor 2022, NE: Normalized Eigenfactor]

| Journal | Preprints | Proportion | JIF | NE |
| --- | --- | --- | --- | --- |
| PLoS One | 1094 | 0.18 | 3.7 | 153.58 |
| Sci Rep | 507 | 0.15 | 4.6 | 227.88 |
| Nat Commun | 437 | 0.42 | 16.6 | 303.23 |
| BMJ Open | 325 | 0.15 | 2.9 | 33.75 |
| Clin Infect Dis | 211 | 0.16 | 11.8 | 29.97 |
| Front Immunol | 192 | 0.08 | 7.3 | 56.67 |
| Elife | 189 | 0.71 | 7.7 | 59.73 |
| Viruses | 173 | 0.09 | 4.7 | 12.83 |
| Front Public Health | 138 | 0.03 | 5.2 | 9.60 |
| Vaccine | 135 | 0.11 | 5.5 | 13.55 |
| J Med Virol | 132 | 0.06 | 12.7 | 7.68 |
| Int J Infect Dis | 129 | 0.10 | 8.4 | 8.68 |
| Proc Natl Acad Sci USA | 129 | 0.21 | 11.1 | 146.31 |
| J Infect Dis | 122 | 0.20 | 6.4 | 12.21 |
| BMC Public Health | 116 | 0.08 | 4.5 | 18.96 |
| PLoS Pathog | 113 | 0.44 | 6.7 | 12.83 |
| PLoS Comput Biol | 112 | 0.54 | 4.3 | 13.61 |
| Microbiol Spectr | 108 | 0.24 | 3.7 | 2.33 |
| JAMA Netw Open | 105 | 0.08 | 13.8 | 35.07 |
| Science | 102 | 0.16 | 56.9 | 172.36 |
| BMC Infect Dis | 101 | 0.1126 | 3.7 | 8.46 |
| Nature | 97 | 0.0968 | 64.8 | 237.43 |
| Sci Total Environ | 94 | 0.0887 | 9.8 | 80.00 |
| Emerg Infect Dis | 91 | 0.1306 | 11.8 | 10.12 |
| BMC Med | 88 | 0.3713 | 9.3 | 8.70 |
| Euro Surveill | 86 | 0.1950 | 19.0 | 7.21 |
| EBioMedicine | 84 | 0.2333 | 11.1 | 9.78 |
| J Clin Microbiol | 83 | 0.3180 | 9.4 | 6.22 |
| Epidemiol Infect | 77 | 0.1897 | 4.2 | 2.40 |
| Cell | 76 | 0.3276 | 64.5 | 102.85 |

## Conclusion and Discussion

This study highlights the significant role of preprints in the dissemination of COVID-19-related research. The publication of 12,152 preprint-derived articles in 1,380 journals, far exceeding the 246 journals accepting direct transfers from bioRxiv, indicates broad acceptance and proactive journal selection by authors. This broad distribution across a wide range of journals reflects the integration of preprints into mainstream scientific communication. High-impact journals tend to show a higher proportion of preprint-derived articles, suggesting their preference among authors and contributing to higher Journal Impact Factor (JIF) and Normalized Eigenfactor scores. Specifically, the top 30 journals, which include prestigious titles such as Nature, Cell, and Science, accounted for approximately 50% of all preprint-derived articles, emphasizing their central role in the rapid dissemination of significant research findings. These findings suggest that preprints are not only facilitating the rapid dissemination of COVID-19-related research but are also being preferentially published in high-impact journals. This underscores the growing acceptance and importance of preprints in the scientific community, contributing to faster communication and higher citation rates.

## Acknowledgments

This work was supported by JSPS KAKENHI Grant Numbers JP19K12707, JP20K12569, JP22K12737, and ROIS NII Open Collaborative Research 2023(23FS01)

## Notes

### Competing Interest Statement

The authors have declared no competing interest.

